# High levels of copy number variation of ampliconic genes across major human Y haplogroups

**DOI:** 10.1101/230342

**Authors:** Danling Ye, Arslan Zaidi, Marta Tomaszkiewicz, Corey Liebowitz, Michael DeGiorgio, Mark D. Shriver, Kateryna D. Makova

## Abstract

Due to its highly repetitive nature, the human male-specific Y chromosome remains understudied. It is important to investigate variation on the Y chromosome to understand its evolution and contribution to phenotypic variation, including infertility. Approximately 20% of the human Y chromosome consists of ampliconic regions which include nine multi-copy gene families. These gene families are expressed exclusively in testes and usually implicated in spermatogenesis. Here, to gain a better understanding of the role of the Y chromosome in human evolution and in determining sexually dimorphic traits, we studied ampliconic gene copy number variation in 100 males representing ten major Y haplogroups world-wide. Copy number was estimated with droplet digital PCR. In contrast to low nucleotide diversity observed on the Y in previous studies, here we show that ampliconic gene copy number diversity is very high. A total of 98 copy-number-based haplotypes were observed among 100 individuals, and haplotypes were sometimes shared by males from very different haplogroups, suggesting homoplasies. The resulting haplotypes did not cluster according to major Y haplogroups. Overall, only three gene families (*DATZ, RBMY, TSPY*) showed significant differences in copy number among major Y haplogroups, and the haplogroup of an individual could not be predicted based on his ampliconic gene copy numbers. Finally, we found a significant correlation between copy number variation and individual’s height (for three gene families), but not between the former and facial masculinity/femininity. Our results suggest rapid evolution of ampliconic gene copy numbers on the human Y, and we discuss its causes.

## Introduction

Studying the Y chromosome provides insights into sex determination, sex-specific disease risks, and evolutionary history that cannot be determined by studying the female genome alone (Skaletsky et al. 2003; van Oven et al. 2013). However, for the vast majority of mammalian species, only female genomes have been sequenced and assembled. Mammalian females have diploid sex chromosomes (XX), which allows easier sequencing and assembly of the X chromosome compared to the highly repetitive haploid Y chromosome (Tomaszkiewicz et al. 2017).

The eutherian sex chromosomes evolved from a pair of autosomes, with the X chromosome keeping the original autosomal size and the Y chromosome shrinking over time. The male-specific region (MSY) constitutes approximately 95% of the length of the Y chromosome. The MSY encompasses a mosaic of euchromatic – X-degenerate, X-transposed, and ampliconic – and heterochromatic sequences. The human MSY is flanked on both sides by pseudoautosomal regions (PARs), the only parts of the Y that recombine with the X (Skaletsky et al. 2003).

The Y chromosome acquired the sex-determining gene, *SRY*, and subsequently underwent a series of inversions that suppressed its ability to recombine with the X chromosome over most of its length (Lahn et al. 2001). As a result, the Y chromosome has become prone to accumulation of deleterious mutations via Muller’s ratchet, genetic hitchhiking along with beneficial alleles, and background selection against deleterious alleles (Charlesworth & Charlesworth 2000; Filatov et al. 2000; Bachtrog 2008, 2013). The Y chromosome is present only in males and is haploid. Therefore, its effective population size is a fraction of that for autosomes, making it more susceptible to genetic drift (Charlesworth & Charlesworth 2000; Filatov et al. 2000). Because the Y is non-recombining over most of its length and inherited exclusively along the paternal lineage, it provides information about patterns of male-specific dispersal and gene flow (Hammer et al. 2008).

Previous studies have noted reduced nucleotide diversity on human MSY relative to autosomes (e.g., Dorit et al. 1995; Wilson Sayres et al. 2014) and attempted to explain this observation by its small effective population size (Charlesworth & Charlesworth 2000; Filatov et al. 2000), high variance in reproductive success among males (Hammer et al. 2008; Wilder et al. 2004), high levels of gene conversion among palindrome arms (Rozen et al. 2003; Marais et al. 2010; Helgason et al. 2015), and purifying selection (Wilson Sayres et al. 2014). In contrast, structural diversity on the Y is known to be high (Repping et al. 2006), which is consistent with frequent intrachromosomal rearrangements facilitated by the repetitive nature of the Y (Skaletsky et al. 2003).

In humans, as in most other mammals studied, the MSY plays an important biological role. It harbors the *SRY* gene that produces the transcription factor initiating male development, while suppressing signals leading to the development of female reproductive organs (Harley et al. 1992). A number of genes located in the MSY are critical to male reproduction, as their deletion can cause spermatogenic failure (Dhanoa et al. 2016). Additionally, the MSY has been implicated in skeletal growth (Tanner et al. 1959), germ-line and somatic tumorigenesis (Kido & Lau 2015), and graft rejection (Kido & Lau 2015; Scott et al. 1997). As the MSY accumulated genes important for male function to resolve sexually antagonistic selection, it is conceivable that some of them are important for the development of sexually dimorphic traits (Dean & Mank 2014; Case & Teuscher 2015).

The human MSY harbors nine multi-copy ampliconic gene families – *BPY, CDY, DAZ, HSFY, PRY, RBMY, TSPY, VCY*, and *XKRY* (Skaletsky et al. 2003; Bhowmick et al. 2007). All but one (*TSPY*) of these gene families are located within either palindromes (P1, P2, P3, P4, P5, and P8) or an inverted repeat (IR2) (Skaletsky et al. 2003). The *TSPY* gene family is arrayed in tandem outside palindromes and more widely spaced inverted repeats (Skaletsky et al. 2003). Seven of the nine families are implicated in spermatogenesis or sperm production, and all nine gene families are expressed predominantly or exclusively in testes (Skaletsky et al. 2003; Bhowmick et al. 2007). Ampliconic gene copies within each family have high sequence identity (>99.9%) that is maintained by gene conversion, which prevents degeneration of these gene families critical for male function (Rozen et al. 2003). It has been proposed that multiple copies of ampliconic genes accumulated on the Y because they increase male reproductive fitness via enhanced sperm production (Rozen et al. 2003; Betrán et al. 2012; Bellott et al. 2014).

Several studies have focused on exploring associations between ampliconic gene copy number and reproductive diseases, and/or fertility. The regions that have been reported to be deleted on the Y chromosome in infertile males are azoospermia factor (AZF) regions a, b and c (AZFa, AZFb, and AZFc), the latter two containing ampliconic gene families (Vogt et al. 1996; Krausz & Degl’Innocenti 2006; Yu et al. 2015). AZFb contains *CDY2, XKRY, HSFY*, and *PRY* families, deletions in which have been shown to lead to spermatogenic arrest (Foresta et al. 2001; Krausz et al. 2014). AZFc contains *DAZ, BPY2, CDY1A*, and *CDY1B* families, deletions in which can result in different levels of spermatogenic failure (Pryor et al. 1998; Krausz et al. 1999) and can be heritable (Page et al. 1999; Rozen et al. 2012). The AZFc region is highly repetitive, harbors palindromes (Kuroda-Kawaguchi et al. 2001) and thus is more prone to deletions than the other AZF regions (Navarro-Costa et al. 2010; Knebel et al. 2011). Indeed, AZFc deletions constitute 80% of all AZF deletions (Bansal et al. 2016). Ampliconic gene families outside of AZF regions are also implicated in reproductive diseases. For example, copy number reductions in *DAZ, BPY*, and *CDY* gene families have been associated with lower total motile sperm counts in men (Noordam et al. 2011; Bansal et al. 2016). Contradictory results have been reported on the association between *TSPY* and fertility (Krausz et al. 2010). Nickkholgh and colleagues (Nickkholgh et al. 2010) did not find a statistically significant difference in *TSPY* copy number between men with low vs. high sperm counts, while Giachini and colleagues (Giachini et al. 2009) reported that low *TSPY* copy number is associated with low sperm production. No studies have been conducted to explore potential associations of Y chromosome ampliconic gene copy numbers and traits besides fertility, e.g. sexually dimorphic traits.

We presently have only limited knowledge about Y chromosome ampliconic gene copy number variation in healthy males within and among human populations. In fact, the only available information comes from the analysis of small samples of persons of European ancestry. Earlier studies have determined copy number for a total of only three males (Tomaszkiewicz et al. 2016; Skaletsky et al. 2003). Recently, Skov and colleagues investigated Y chromosome ampliconic gene copy number variation in 62 males of Danish descent (Skov et al. 2017).

In the present study, we experimentally determined the copy number of all nine ampliconic genes in 100 men representing ten major Y haplogroups (Y Chromosome Consortium 2002) using droplet digital PCR (ddPCR) (Hindson et al. 2011; McDermott et al. 2013). We used these data to obtain a view of ampliconic gene copy number variation within and across human populations around the world by addressing the following questions: (i) Are ampliconic genes more variable between major Y haplogroups than within haplogroups? (ii) Can ampliconic gene copy number variation be used to classify major Y haplogroups accurately? (iii) How variable are haplotypes reconstructed based on ampliconic gene copy number? (iv) Does ampliconic gene copy number variation underlie variation in sexually dimorphic traits such as height and facial masculinity/femininity? Thus, by answering these questions, we characterized evolution of ampliconic gene copy number variation in a large number of individuals representing major Y haplogroups.

## Materials and Methods

### Sample collection, consent, SNP typing, and DNA extraction

A total of 100 men were recruited with written informed consent as part of the ADAPT and ADAPT2 studies (IRB #44929 and #45727) conducted at the Pennsylvania State University. According to the approved protocol, saliva samples were obtained and two phenotypes – height and facial masculinity/femininity (see below) – were measured for all participants. The saliva samples were sent to 23andMe for genotyping on their v3 and v4 arrays (23andMe, Mountainview, CA). DNA was extracted from the saliva samples using a salting-out method followed by an ammonium acetate cleanup (Quinque et al. 2006) and quantified using Qubit dsDNA BR Assay Kit (Invitrogen, Carlsbad, CA).

### Droplet digital PCR (ddPCR)

For each of the 100 DNA samples, we performed ddPCR for nine ampliconic gene families of interest (*BPY, CDY, DAZ, HSFY, PRY, RBMY, TSPY, VCY*, and *XKRY*) and for *SRY*, a single-copy gene on the Y chromosome, used as a reference. Each sample was run in triplicates. In 24 cases (out of a total of 900) one replicate had no calls, and in one case two replicates had no calls (Table S1A). The ddPCR copy number assays were performed using the QX200 system and EvaGreen dsDNA dye (Bio-Rad, Hercules, CA) using the protocol and primers described in our previous publication (Tomaszkiewicz et al. 2016). Briefly, for a completion of one assay replicate for each DNA sample included in the study, *BPY, CDY, HSFY, TSPY*, and *XKRY* were amplified at an annealing temperature of 59°C on one plate, and *DAZ, PRY, RBMY* and *VCY* were amplified with an annealing temperature of 63°C on another plate. *SRY* was amplified on each plate for the ampliconic gene copy number inference. Based on the human reference genome sequence, the primers designed were specific for capturing functional ampliconic gene families (one primer pair per gene family) except for *TSPY*, for which primers were designed to anneal to the smallest number of pseudogenes (Tomaszkiewicz et al. 2016).

The fluorescence in each droplet was measured and an automatic threshold was drawn using QuantaSoft software (Bio-Rad, Hercules, CA). Droplets above the threshold were counted as positive, and those below it were counted as negative. The concentration (copies/uL) of the ampliconic gene family of interest was divided by the concentration of the reference, *SRY*, a single-copy gene in a human male genome (Tomaszkiewicz et al. 2016). Because each sample was run in triplicates, we had three measurements (or two measurements when one of the replicates had no call) of ampliconic gene copy number for each gene family in every individual. Where three replicates were present, the observation most distant from the median was removed to reduce the effect of outliers (Table S1A). After this, ampliconic gene copy number was determined by calculating the mean across the two replicates for each sample (Table 1B). We present the median, standard deviation (SD) and coefficient of variation across individuals for each gene family in Table 2 and Fig. S1.

**Table 1.** Male samples utilized in the study.

| Major Y haplogroups | Y sub-haplogroups | Number of males |  | Major geographic location (Karmin et al. 2015) |
| --- | --- | --- | --- | --- |
| C | C3 | 5 | 5 | Asia |
| E | E1b1a | 5 | 22 | Africa |
|  | E1b1a1a1g1a | 7 |  |  |
|  | E1b1b1 | 5 |  |  |
|  | E1b1b1a | 5 |  |  |
| G | G2 | 5 | 5 | Africa |
| I | I1 | 5 | 15 | Europe |
|  | I2a2a | 5 |  |  |
|  | I2Aa1b | 5 |  |  |
| J | J2 | 5 | 5 | Western Asia |
| L | L1 | 4 | 4 | Western Asia |
| O | O1 | 3 | 14 | Eastern and Southeastern Asia |
|  | O2 | 6 |  |  |
|  | O3 | 5 |  |  |
| Q | Q1 | 5 | 5 | Central Asia |
| R | R1b1a2a1a2c | 5 | 20 | Europe |
|  | R1b1a2a1a2b | 5 |  |  |
|  | R1b1a2a1a1 | 5 |  |  |
|  | R1a1a1 | 5 |  |  |
| T | T | 5 | 5 | Western Asia |
| Total |  | 100 |  |  |

**Table 2.**
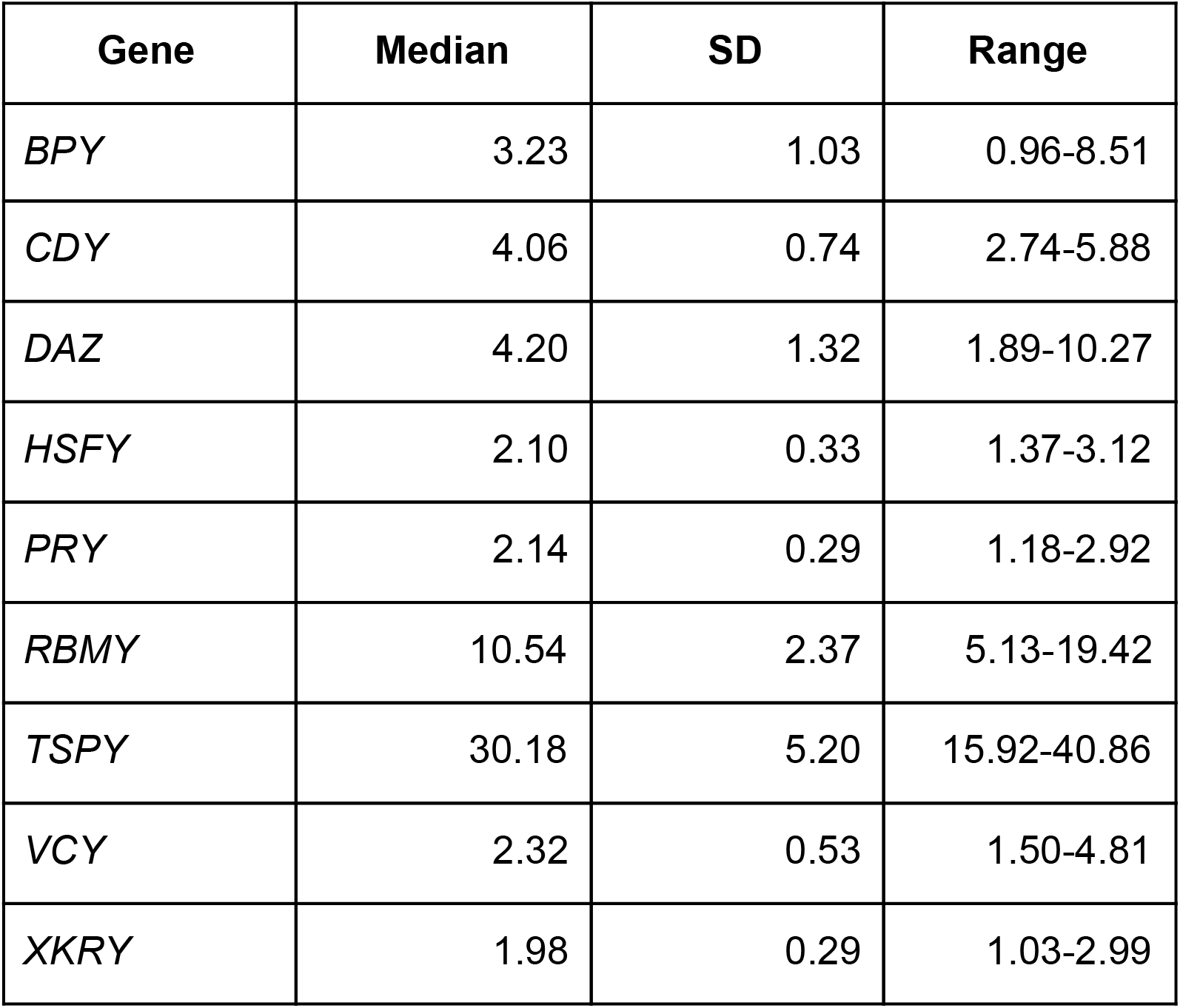
Median, standard deviation (SD) and range of unrounded copy number values per ampliconic gene family (based on the data from Table S1A).

### Construction of phylogeny based on SNP data

A maximum likelihood phylogenetic tree based on 187 segregating Y chromosome SNPs for 100 male individuals was constructed based on the Tamura-Nei model using MEGA7 (Kumar et al. 2016). The initial trees for the heuristic search were obtained automatically by applying the BioNJ algorithm (Gascuel 1997) to a matrix of pairwise distances estimated using the Maximum Composite Likelihood (MCL) approach, and then selecting the topology with the highest log likelihood value.

### Evaluating differences in ampliconic gene copy numbers among haplogroups

We tested whether ampliconic gene copy number is different among different haplogroups for each gene family separately. This was done using two different approaches. First, we applied the conventional one-way analysis of variance (ANOVA), which does not take into account the phylogenetic relationships among Y-haplogroup lineages. The simple ANOVA was performed for each ampliconic gene family using major haplogroup (C, E, G, I, J, L, O, Q, R, and T) as factor.

Second, we applied the EVE model (Rohlfs & Nielsen 2015), which accounts for the phylogenetic structure among haplogroups. Whereas the EVE model was developed with the intention of testing for non-neutral evolution of gene expression in a given phylogeny, it can be applied to any quantitative trait as long as it is measured on multiple individuals from every species in the phylogeny (Rohlfs & Nielsen 2015). Our goal was to measure the ratio of variation in copy number within haplogroups to the variation between haplogroups, denoted by *β_i_* for every gene family, i = 1,2,…, 9. We expect this ratio to be similar across gene families evolving neutrally in the phylogeny (i.e. *β_i_* = *β_shared_*, i = 1,2,…, 9). Deviations from this expectation can be suggestive of selection. As such, we test whether *β_i_* for any one gene family *i* deviates from this expectation (i.e *β_i_* ≠ *β_shared_*). If *β_i_* < *β_shared_*, then there is more variation across haplogroups than the variation within haplogroups, which could be suggestive of directional selection in some haplogroups. Conversely, if *β_i_* > *β_shared_*., then there is more variation within haplogroups than variation across haplogroups, which could be indicative of high conservation of copy number across haplogroups.

To apply the EVE model to the copy number data, we first constructed an ultrametric tree connecting the major haplogroups from the phylogenetic tree based on Y-chromosomal SNPs. This was done by first collapsing all individual branches from the same haplogroup such that each major haplogroup is represented by one terminal branch in the phylogeny. Then, we scaled the tree by setting the time of the most recent common ancestor of all lineages to 72,500 years ago based on the most recent common ancestor (MRCA) of the major haplogroup lineages represented in our dataset and the Y phylogeny presented by Karmin and colleagues (Karmin et al. 2015). We estimated the parameter *β_i_* for each gene from the copy number data using EVE, as well as the *β_shared_* across all genes, and calculated the likelihood ratio between the null hypothesis (*H*_0_: *β_i_* = *β_shared_*) and alternative hypothesis (*H*_1_ :*β_i_* ≠ *β_shared_*). A P value for each test was calculated assuming that the likelihood ratio asymptotically follows a chi-square distribution with one degree of freedom. The likelihood ratio for each gene and corresponding values are presented in Table 3.

**Table 3.**
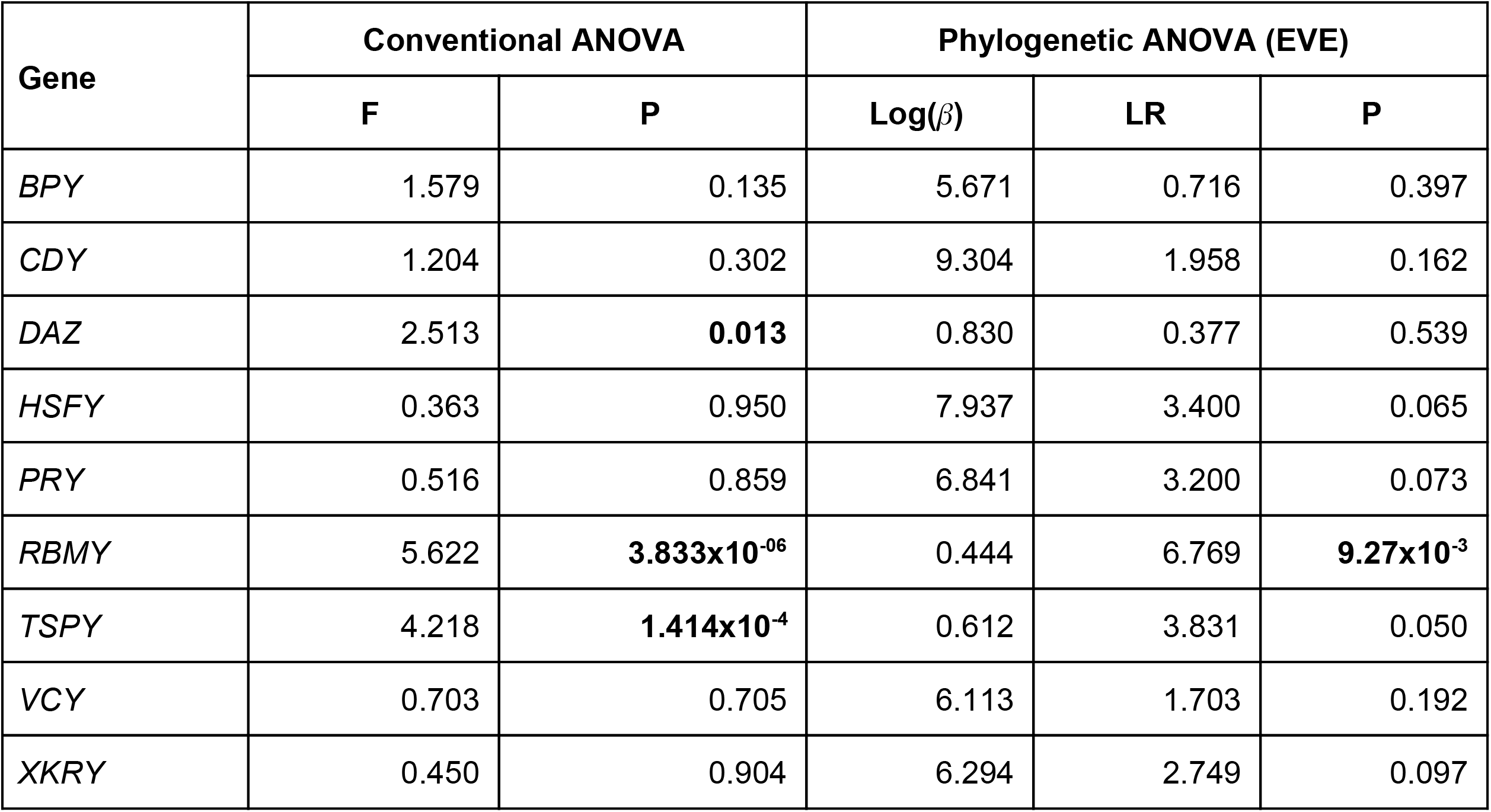
Analysis of variance of the ampliconic gene copy number data. Both conventional one-way ANOVA and phylogenetic ANOVA (EVE) were performed to determine which ampliconic gene families vary significantly in their copy numbers among major haplogroups. F is the f-statistic for the one-way ANOVA. *β* and LR are the ratio of the within-haplogroup variance to the between-haplogroup variance in copy number and the likelihood ratio between the null model and the alternative model, respectively, from the phylogenetic ANOVA (see Methods).

### Clustering of major haplogroups by copy number

Principal Component Analysis (PCA) was performed on the centered and scaled ampliconic gene copy numbers 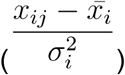, where *x_ij_* is the copy number of the i^th^ gene family and j^th^ individual, to visualize the clustering of major haplogroups based on ampliconic gene copy number (Cirillo 2016). For comparison, we also carried out PCA on the genotypes of SNPs on the Y chromosome using Plink 1.9 (Chang et al. 2015).

In addition to the unsupervised PCA, we also carried out Linear Discriminant Analysis (LDA) to determine whether ampliconic gene copy number of an individual can be used to correctly predict their major haplogroup. This was carried out using the lda function in the MASS package in R (Venables & Ripley 2002). With leave-one-out cross validation, we calculated the posterior probability that each individual can be assigned to their correct haplogroup (Fig. 5).

### Haplotype variability and network analysis

Rounding the fractional copy numbers generated by ddPCR could artificially introduce variation in the data, which could overestimate the number of haplotypes. To evaluate whether this was the case, we calculated the range of haplotypes observed by randomly rounding the original data – the values produced by averaging the two most similar replicates for each gene family and individual – up or down (i.e. floor or ceiling) (Tables S1B and S4A). This was done by generating 100 sets of haplotypes, each of which was obtained by rounding a value *y* either up or down if [floor(y) + 0.25] < *y* < [ceiling(y) – 0.25] where floor(y) refers to the greatest integer less than *y* and ceiling(y) refers to the smallest integer greater than y. Values outside this range were rounded to the nearest integer. For example, a mean copy number of 2.35 was either rounded up or down to 2 or 3, respectively, but a copy number of 2.15 was always rounded down to 2. We performed the same experiment on unrounded ampliconic gene copy numbers from the data in Skov et al. (Table S3A) (Skov et al. 2017). A total of 100 data sets, each consisting of randomly rounded values for each of the 100 (our data set) and 62 (Skov et al.’s data set) individuals, were produced (Table S4A and S4B) and the range of the number of haplotypes observed was calculated (Table S5B, Fig. S3). We found the number of haplotypes in our data set to vary form 98 to 100 (median = 99, Table S5A) and in the Skov data set to vary from 40 to 52 (median = 45; Table S5B).

Haplotype networks based on Y-chromosomal SNP genotypes and based on ampliconic gene copy numbers were constructed separately. The alignment of SNP genotypes from 100 males was inserted as an input for reconstructing haplotypes using “pegas” package in R (Paradis 2010; Cirillo 2016). To construct haplotype networks, we rounded the copy numbers to the nearest integer for both our and Skov et al.’s (Skov et al. 2017) data sets. The alignment of nine different ampliconic gene copy numbers from each of the 100 male individuals was used to build a haplotype network accounting specifically for indel mutations using “haplotypes” package in R (Cirillo 2016). The same approach was used to construct the haplotype network for 62 males from the Danish population (Skov et al. 2017). Haplotype distance matrices used for the haplotype network reconstructions are provided in Tables S4A and S4B. Haplotypes were separated by deletions or insertions of ampliconic gene copies, and each link reflected one-copy number difference. For instance, two haplotypes differing only by two copies of *TSPY* (and having the same copy numbers for the other gene families), 18 and 20, were separated by two links. Similarly, two haplotypes, differing in copy number of two gene families, e.g., *TSPY* and *RBMY*, by one copy each (in the first haplotype *TSPY* = 18 and *RMBY* = 10, while in the second haplotype *TSPY* = 19 and *RMBY* = 9) were also separated by two links.

To get an idea of which ampliconic gene families were contributing most to the variability observed among haplotypes, we sampled pairs of haplotypes, uniformly at random, separately from within and between major Y haplogroups, and counted the copy number differences per ampliconic gene family between each pair. A total of 1,000 such pairs for each comparison, within and between major haplogroups, were generated. The results are shown in Fig. 8.

### Measurement of height and facial masculinity/femininity (FMF)

For the participants in the ADAPT study (a total of 64 men), height was measured using a standard stadiometer. Self-reported height was used for 36 participants from the ADAPT2 study due to remote sampling and lack of a portable stadiometer. Facial masculinity was calculated from 3D images collected on participants using a method developed by (Claes et al. 2014), as described briefly below. FMF scores were estimated by orthogonally projecting the participants’ faces onto the regression line that represents facial sexual dimorphism. A spatially dense mesh of 7,150 quasi-landmarks (QL) was superimposed on participant’s 3D facial scans and differences in translation, rotation, and scale were removed by applying a Generalized Procrustes Superimposition (GPS) on the set of facial coordinates (Claes et al. 2014). The first sixty principal components, which explained 98% of the variance, were retained. To calculate FMF, we used a leave-one-out cross-validation approach, that is, the participant face for whom we wanted FMF to be estimated was left out of the regression model while the remaining participants were used to estimate regression coefficients with a multivariate linear regression of facial Principal Components on sex and height. Height was used too as a covariate to remove the influence of size differences on facial shape from the estimation of FMF. The average female face was set as the origin of the facial PCA, allowing higher values to reflect more masculine faces. Using the regression line for sex, the FMF score was orthogonally projected for the participant’s face. Both height and FMF data are provided in Table S6.

### Evaluating correlations between haplogroups and phenotypic traits

We evaluated correlations between ampliconic gene families and phenotypic traits using the phylogenetic generalized least square method (PGLM) implemented using the nlme package in R (Cirillo 2016; Pinheiro J, Bates D, DebRoy S, Sarkar D and R Core Team 2017). As some individuals were more closely related to each other than to other individuals, the ampliconic gene copy number data for each individual cannot be considered to be independent data points. To take this phylogenetic relatedness into account, we constructed a variance-covariance matrix from the ultrametric Y-chromosomal phylogeny using the vcv function in the ape package in R (Paradis et al. 2004), assuming a Brownian motion model of phenotypic evolution (Wilson Sayres et al. 2011; Cirillo 2016). This variance-covariance matrix was used to specify the correlation structure of the residuals.

We tested whether ampliconic gene copy number for each of the nine ampliconic gene families is a predictor of the two phenotypic traits using the gls function from the nlme package in R (Cirillo 2016). The models were fit using maximum likelihood and significance of the ampliconic gene copy number as a predictor of height, and FMF was determined using a likelihood ratio test between the “full” (intercept + predictor) and “reduced” (intercept only) models.

### Code availability

All the scripts for this study are provided at GitHub: https://github.com/makovalab-psu/Ampliconic_CNV

## Results

### Ampliconic gene copy number variation

To study copy number variation of Y chromosome ampliconic genes, we applied ddPCR. This method allows absolute quantification of the target DNA copies without the need to run a standard curve. This is in contrast to other methods such as quantitative real-time PCR (qRT-PCR), in which suboptimal amplification efficiency influences cycle threshold values and can ultimately result in an inaccurate quantification of the target (Hindson et al. 2011; McDermott et al. 2013; Pinheiro et al. 2012). ddPCR was recently used to evaluate the copy number of ampliconic Y chromosome genes in humans and gorillas (Tomaszkiewicz et al. 2016) and to verify computationally derived ampliconic gene copy number estimates for chimpanzees and bonobos (Oetjens et al. 2016).

In this study, the ddPCR assay, with the primers previously developed by us (Tomaszkiewicz et al. 2016), was used to estimate the copy number for Y chromosome ampliconic genes in 100 male participants from the ongoing Anthropometrics, DNA and the Appearances and Perceptions of Traits (ADAPT) study. The goal of the ADAPT study (http://ched.la.psu.edu/projects/adapt), based at the Pennsylvania State University, is to study the evolutionary, genetic, and socio-cultural factors shaping complex phenotypic variation within and across human populations. Among ADAPT participants, we selected 100 males harboring Y chromosomes from ten major haplogroups (Y Chromosome Consortium 2002): C, E, G, I, J, L, O, Q, R, and T (Table 1). Individuals with subhaplogroups that are evolutionarily close to each other were grouped into a ‘major haplogroup’ category to increase the statistical power in subsequent analyses. For example, individuals from the O1, O2, and O3 subhaplogroups were grouped into the ‘O’ major haplogroup category. These haplogroups were selected because they find their origins in different regions of the world (Table 1).

The copy number for each gene family for every individual was estimated using three technical replicates, with a handful of exceptions for which fewer than three replicates were analyzed (Table S1A). In total, we processed 100 males x 9 gene families = 900 samples, 875 of which were analyzed in three replicates. To assess the consistency of measurements among replicates, we calculated the coefficient of variation (i.e. standard deviation divided by mean), CV, across replicates. The median CV was low, 3.5% of the mean across all samples (red dashed line in Fig. S1A). After removing the most distant value among the three replicates (see Methods), the median CV was even lower; 1.07% of the mean (red dashed line in Fig. S1B). We averaged the values of the two remaining replicates and used them in all subsequent analyses (Tables S1A and S1B). We used these unrounded average values for all the analyses, except for counting the number of haplotypes and building haplotype networks, where we rounded the averaged values to the nearest integer.

### Variation in copy number among gene families

We first tested whether larger gene families were also more variable in their copy number among individuals. Such a relationship is expected because the probability of copy insertions and deletions increases with copy number (Ghenu et al. 2016). Indeed, the median copy number for ampliconic gene families across individuals is positively correlated with the variance in copy number (Spearman’s r = 0.99; Fig. 1). Larger gene families are indeed more variable, on average (Fig. 1; Table 2).

**Figure 1.**
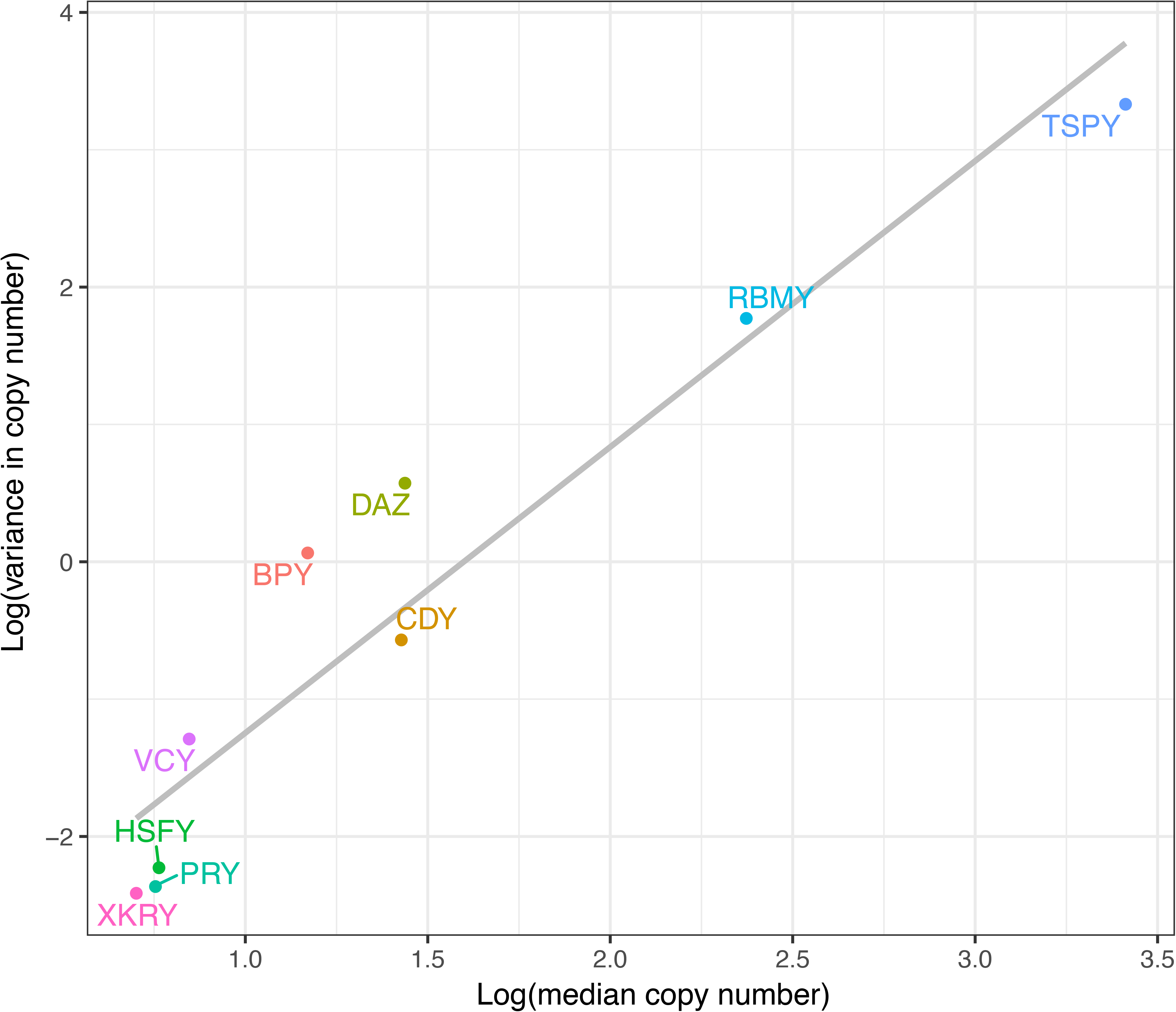
Larger gene families tend to be more variable. The median and variance of copy number were calculated across all individuals in the sample (N = 100). The grey line shows the line of best fit (from ordinary least squares regression).

### Lack of a phylogenetic pattern in ampliconic gene copy number variation

To examine whether there is a phylogenetic pattern underlying ampliconic gene copy number variation in the humans studied, we constructed a phylogenetic tree based on Y chromosome single nucleotide polymorphisms (SNPs) and superimposed copy numbers for each of the ampliconic gene families per individual next to this phylogeny (Fig. 2), following (Skov et al. 2017). As expected, individuals from the same haplogroup clustered together based on Y chromosome SNPs. However, ampliconic gene copy number variation did not show discernible patterns with respect to the Y-specific phylogeny.

**Figure 2.**
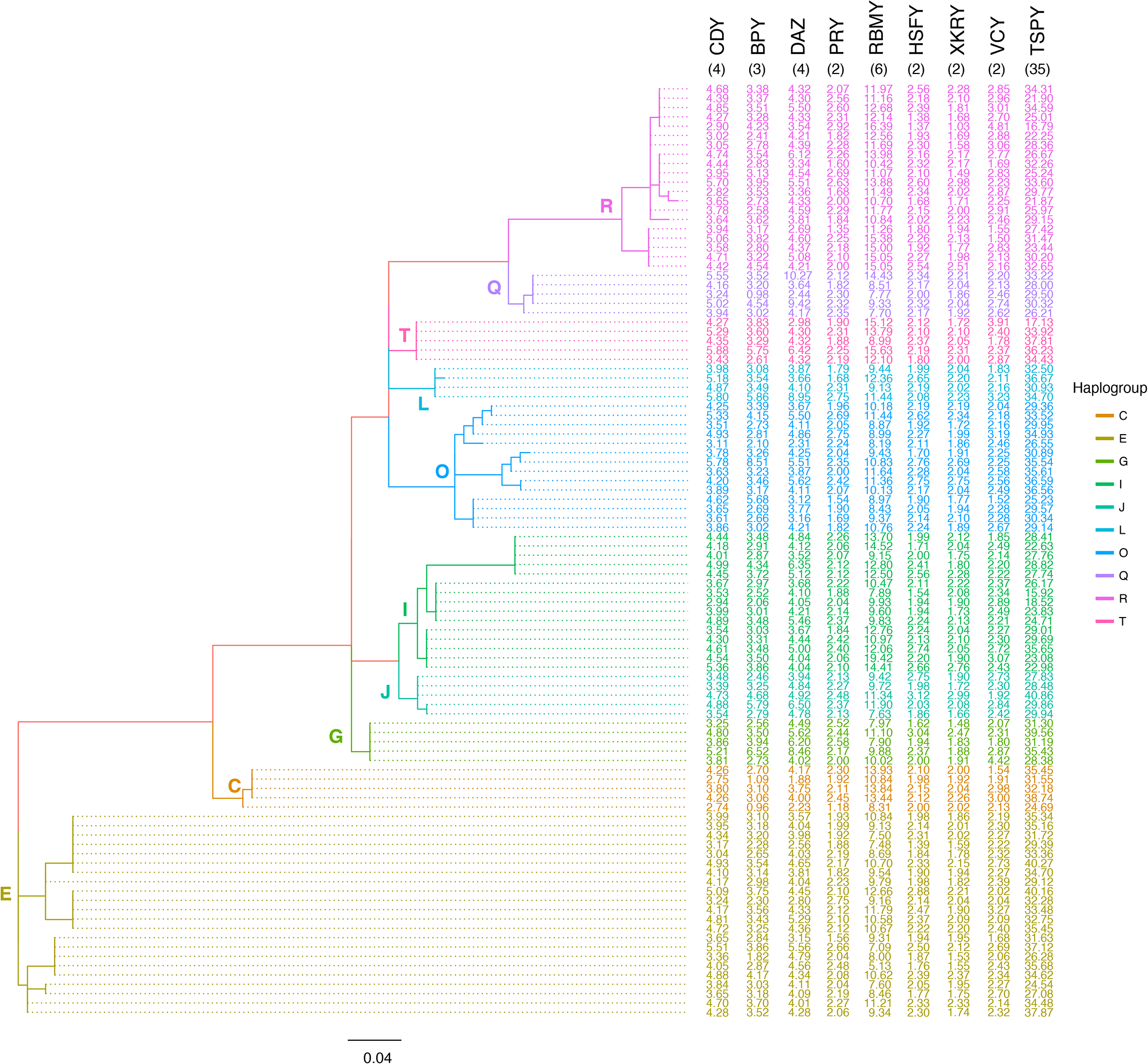
The phylogenetic tree based on Y-chromosomal SNPs. The evolutionary tree was inferred from 187 Y-chromosomal SNPs using maximum likelihood (log-likelihood = −993.63). The branches are colored according to Y haplogroup. Ampliconic gene copy number averaged between two most similar replicates is presented on the right. For comparison, we included the copy numbers for an individual sequenced by Skaletsky and colleagues (indicated in black font in parenthesis) (Skaletsky et al. 2003).

### Differences in ampliconic gene copy numbers among Y haplogroups

We further tested whether ampliconic gene copy numbers are significantly different among the ten major Y haplogroups analyzed. The distribution of ampliconic gene copy numbers per family across all Y-haplogroups is shown in Figure 3. Using a one-way ANOVA test (Table 3) we found that copy numbers of *BPY, CDY, HSFY, PRY, VCY*, and *XKRY* gene families were not significantly different among major Y haplogroups. However, copy numbers for *DAZ* (P = 0.013), *RBMY* (P = 3.833 x 10^−06^) and *TSPY* (P = 1.414 x 10^−04^) did differ significantly among major haplogroups (Table 3). The differences for the *DAZ* gene family were not significant after Bonferroni correction for multiple tests.

**Figure 3.**
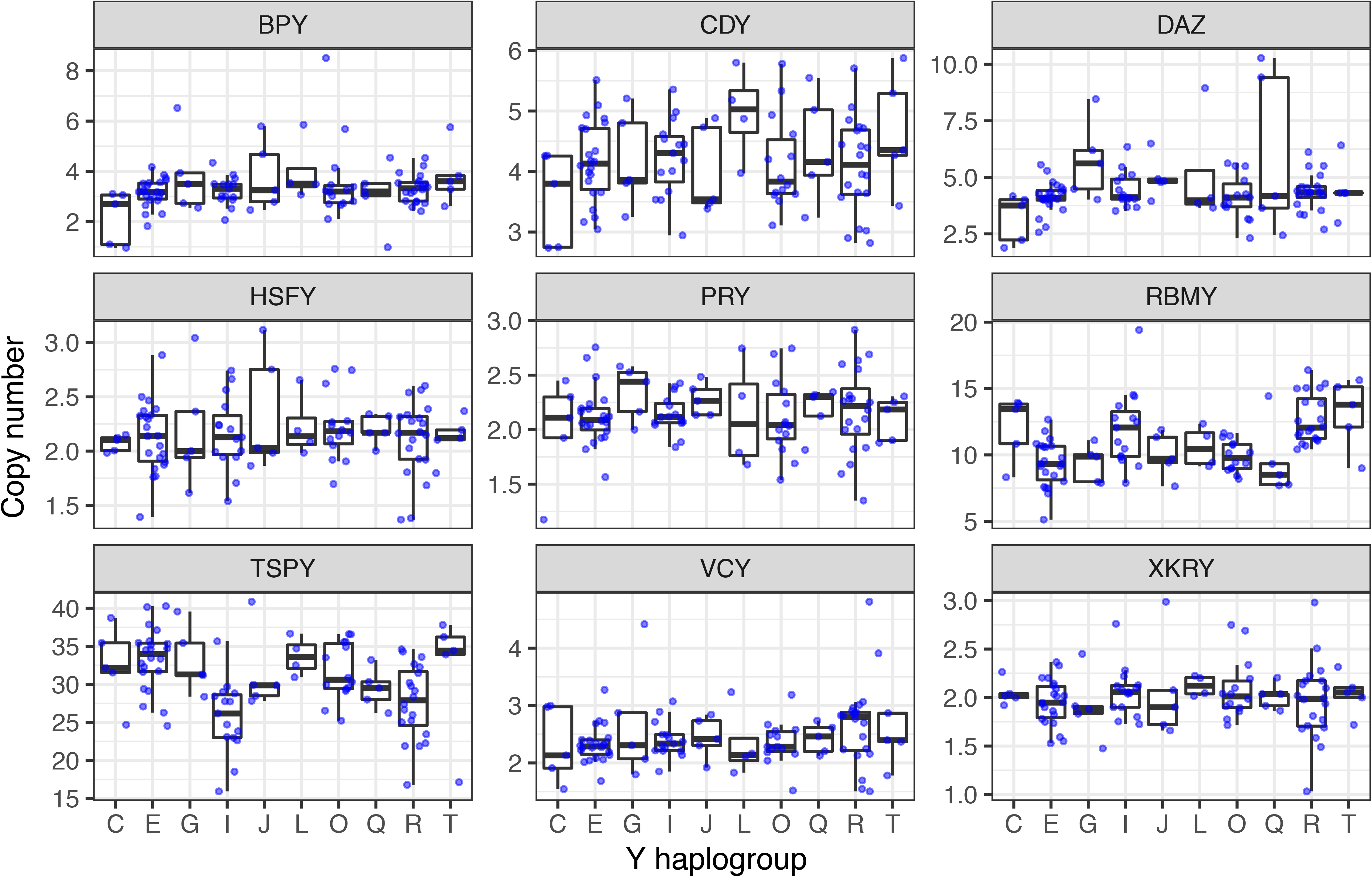
The distribution of ampliconic gene copy numbers across major Y haplogroups. Between four and 22 individuals per major Y-haplogroup were analyzed (see Table 1 for sample sizes for each haplogroup).

In addition to the conventional, one-way ANOVA, we carried out a phylogenetic ANOVA with the Expression Variance and Evolution (EVE) model (Rohlfs & Nielsen 2015). The test estimates a parameter for each gene *i*, *β_i_*, which is the ratio of the variance in ampliconic gene copy number within haplogroups to the variance between haplogroups. It assumes that genes sharing their variability level will share a common *β* parameter, *β_shared_*. Based on a likelihood ratio test, we used EVE to identify genes with either *β_i_* < *β_shared_* (higher variation between haplogroups than within haplogroups), or *β_i_* > *β_shared_* (higher variation within haplogroups than between haplogroups). We found that *RBMY* (log_10_ *β* = 0.444, LR = 6.769, P = 9.270 x 10^−03^) exhibited significantly lower values of *β_i_* than of *β_shared_* (log_10_ *β_shared_* = 1.201; Table 3). *TSPY* (log_10_ *β* = 0.612, LR = 3.831, P = 0.050) also showed a lower value of *β_i_* than of *β_shared_*, which was marginally significant. This result suggests that these two gene families might have diverged more across haplogroups than the overall level of divergence observed in all gene families together. Such cases suggest non-neutral evolution along the phylogeny.

### Do major haplogroups cluster by ampliconic gene copy number?

Because copy numbers for some ampliconic gene families are significantly different among major haplogroups (Table 3), we next tested whether individuals cluster based on ampliconic gene copy number. To answer this question, we carried out Principal Component Analysis (PCA) on ampliconic gene copy numbers. The first three PCs explain ~70% of the total variation (Fig. S2A). The resulting clustering of individuals indicated that, whereas there is some separation of major haplogroups based on ampliconic gene copy number (Fig. 4A-B), it is not nearly as pronounced as clustering based on Y chromosome SNPs (Fig. 4C-D; Fig. S2B).

**Figure 4.**
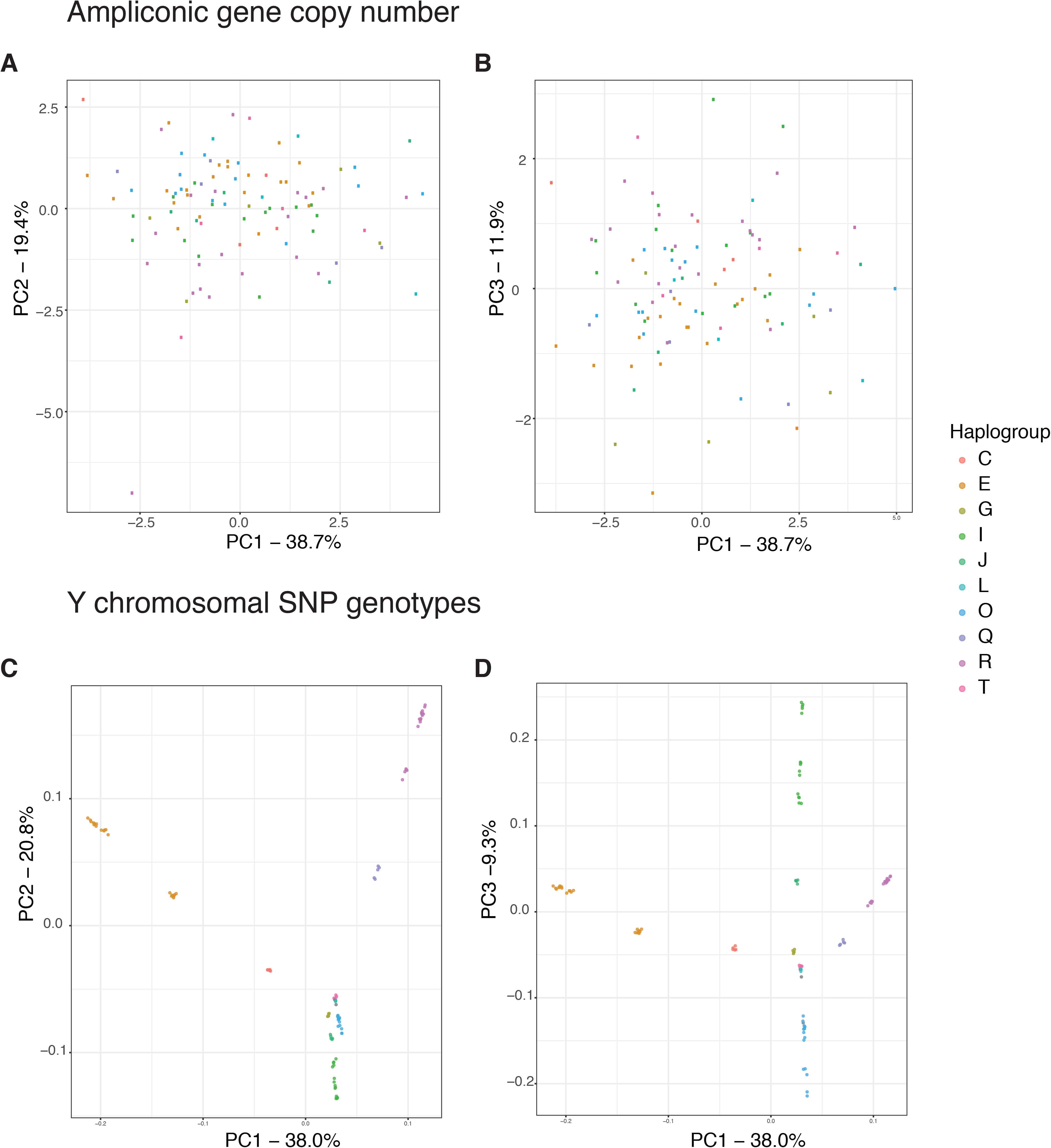
(**A**) and (**B**) Results of PCA on ampliconic gene copy number data (A. PC1 vs PC2; B. PC1 vs PC3). (**C**) and (**D**) Results of PCA on SNP genotype data (C. PC1 vs PC2, D. PC1 vs PC3). Individuals are colored based on the haplogroup determined from SNP genotype data. Individuals cluster by haplogroup based on SNP genotype data but not clearly based on ampliconic gene copy number.

### Can an individual’s haplogroup be predicted based on ampliconic gene copy number?

To test whether we can correctly classify the haplogroup of an individual based on his ampliconic gene copy numbers, we carried out Linear Discriminant Analysis (LDA) with major haplogroup as the response variable and all nine ampliconic gene copy numbers as predictors. Using a leave-one-out approach, we determined the posterior probability that an individual belongs to a major haplogroup based on his copy number profile. The results are displayed as barplots in Figure 5, where individuals are represented by a pair of vertical bars and the probability of being classified correctly (blue), or incorrectly (orange), in the known haplogroup (determined by SNPs) is represented by the height of the bars. We can conclude that the major haplogroups are often ambiguously or incorrectly predicted from copy number variation data alone, which confirms the patterns seen in the PCA plots (Fig. 4), i.e. that most of the variation in ampliconic gene copy number is shared among haplogroups. Consequently, it is difficult to predict the haplogroup of a person based on his ampliconic gene copy number profile.

**Figure 5.**
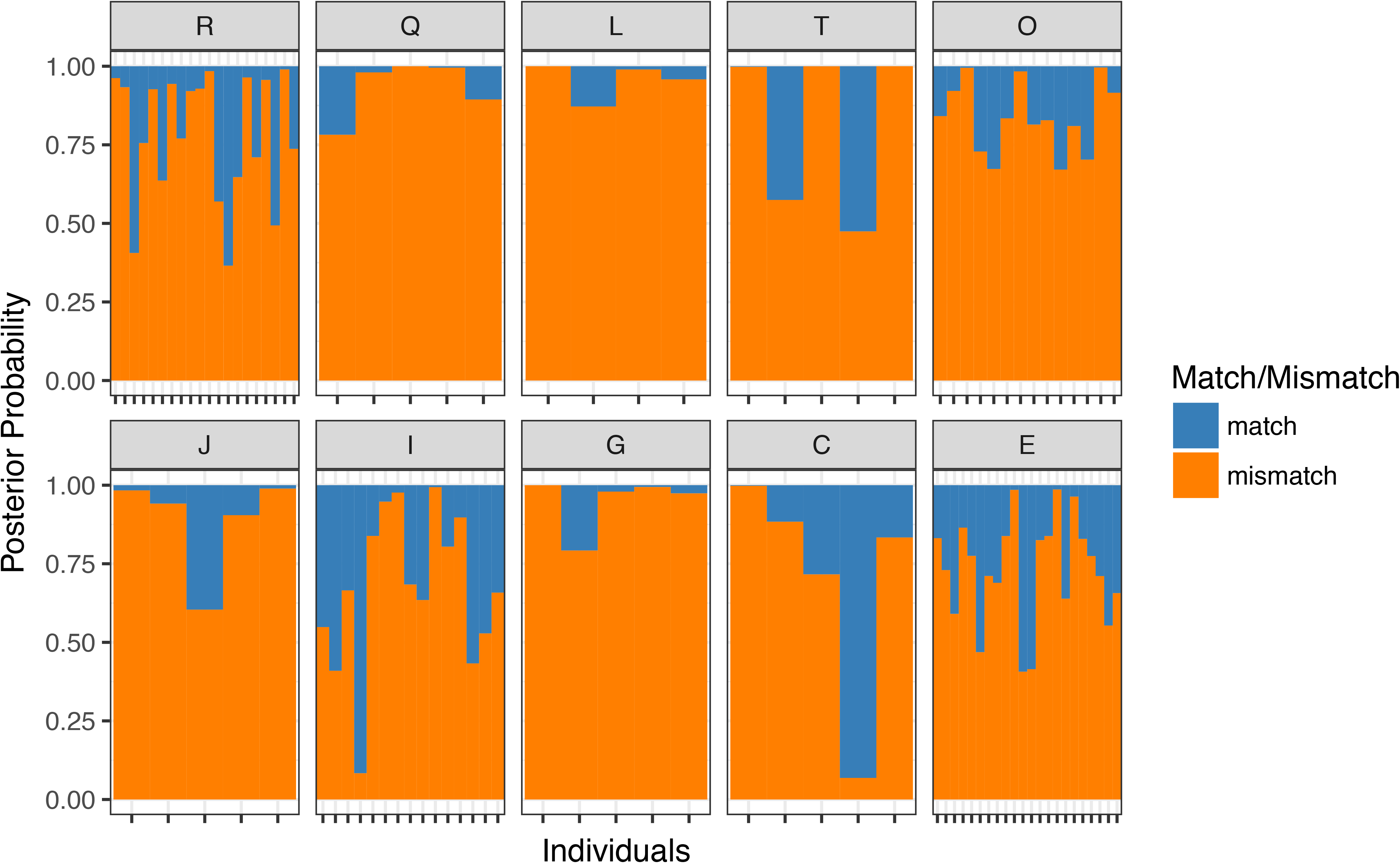
Barplots showing the posterior probability of classifying each individual to his known haplogroup correctly (blue) vs. incorrectly (orange). The known haplogroup of the individual, determined by SNP genotypes, is written on top of each bar plot in the strip.

### Haplotype variability and network analysis

We next compared the variability of haplotypes based on SNP data versus that based on ampliconic gene copy numbers. Based on 187 SNPs on the Y chromosome (from a total of 450 Y-chromosomal SNPs analyzed), there are 39 distinct haplotypes among 100 individuals that cluster, as expected, by either subhaplogroup or major haplogroup (Fig. 6). In fact, many haplogroups are monophyletic, and usually a unique substitution path leads to each haplotype.

**Figure 6.**
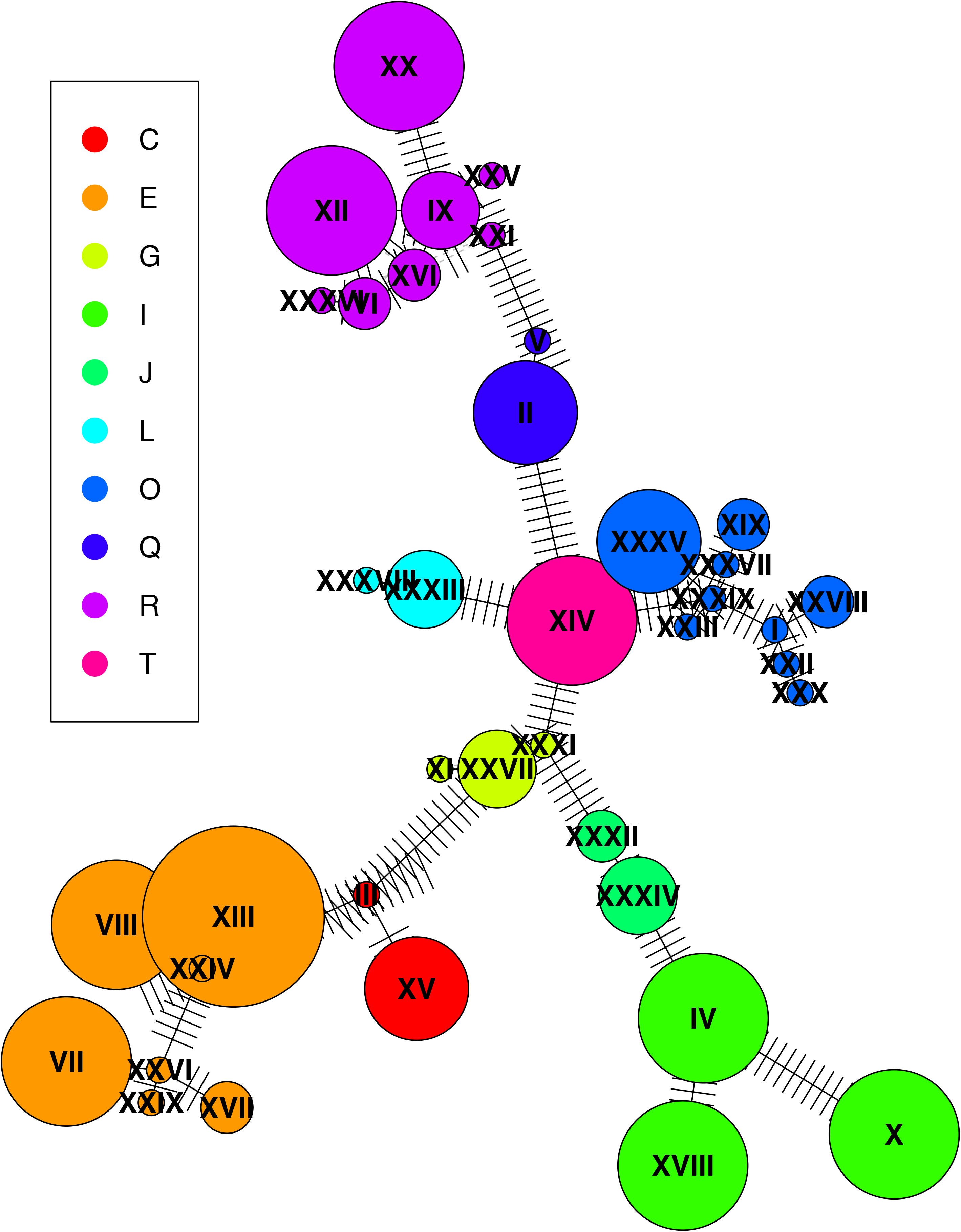
Haplotype network constructed based on Y SNP genotypes from 100 males (39 haplotypes). The disc size is proportional to the number of individuals with a particular haplotype. Black lines connect each haplotype to its closest haplotype, while perpendicular bars correspond to mutational steps between connected haplotypes.

For the same 100 individuals, haplotypes obtained from ampliconic gene copy numbers were more numerous than those obtained from SNP data. To construct haplotypes using ampliconic gene copy numbers, we rounded the values we obtained with ddPCR (after averaging of the two most similar replicates) to the nearest integer (Table S1C). This resulted in 98 haplotypes among 100 individuals studied, more than twice the number of haplotypes obtained from SNP data (Table S2A). The large number of haplotypes observed with copy number data was not because of variation introduced by rounding to the nearest integer (see Methods). The 98 distinct haplotypes usually differed from each other by several copies of genes either from the same or different families (Table S2B). From a total of 4,753 pairwise comparisons among haplotypes, only 64 pairs (~1%) showed a one-copy difference in one gene family (Table S2B). Among the two shared haplotype pairs observed in our sample of 100 males, one pair included a male with an African (E) and a male with an Asian (O2) haplogroups, whereas in the other pair, one male had a European (I) and another one an Asian (Q) haplogroup (Table S2A). Thus, shared haplotypes in these instances provide examples of homoplasy. In a summary, nine ampliconic gene families still produced a greater number of haplotypes than 187 SNPs.

We also studied the variability of ampliconic gene copy number-based haplotypes using rounded ampliconic gene copy number from the data set generated by Skov and colleagues (Skov et al. 2017) (Table S3A). Even though their data set includes 62 Danish males representing only three major European haplogroups (I, R, and Q; Fig. 7B), we observed a total of 35 copy number-based haplotypes (Table S3B), including 22 haplotypes carried by one individual each, and 13 haplotypes shared by two or more individuals. This network (Fig. 7B) displayed more reticulations than the one based on our data (Fig. 7A). One-copy differences within the same ampliconic gene family constituted a small proportion of haplotype pairwise comparisons (16%, 97 from a total of 595 haplotype pairwise comparisons; Table S3C). This proportion was higher than in our data (16% vs. 1%) likely because Skov and colleagues (Skov et al. 2017) only analyzed individuals of Danish ancestry, while we analyzed a world-wide sample. Again, several cases of homoplasy were observed (Table S3B), including the same haplotypes carried by individuals belonging to different major Y haplogroups. Therefore, independently of the divergence time of the studied individuals – worldwide human populations vs. a single Danish population – the number of haplotypes based on ampliconic gene copy number was high. Furthermore, in contrast to the SNP-based haplotype network, the haplotype networks constructed using ampliconic gene copy numbers from the same 100 individuals did not display clustering by major Y haplogroups for both our and Skov *et al.’s* data sets (Fig. 7A-B).

**Figure 7.**
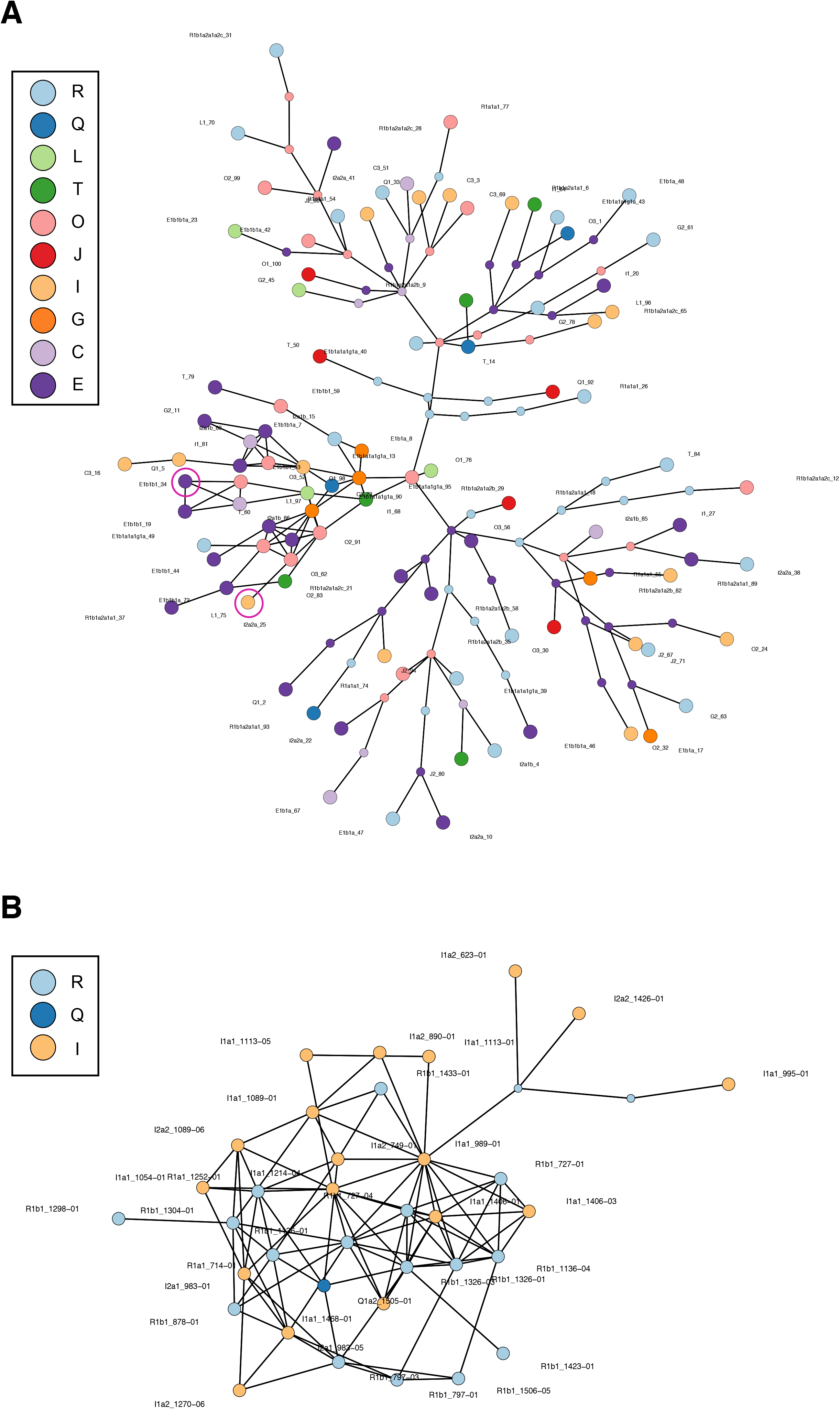
(**A**) Haplotype network constructed based on nine different ampliconic gene copy numbers (rounded) from each of the 100 male individuals (98 haplotypes; rounded copy number values were used; Table S1C). Each big colored disc represents a different haplotype. Small colored discs represent intermediate haplotypes. Black lines connect each haplotype to its closest relative. A link between two haplotypes corresponds to a one-copy difference in one gene family. If extant or ancestral haplotypes are joint by several consecutive links, this indicates several copy number differences (either within the same or different gene families) between them, and the number of such links corresponds to the number of copy number differences. Pink rings indicate haplotypes that were observed in more than one individual. (**B**) Same as A, but for the data from 62 Danish males in (Skov et al. 2017) (rounded copy number values were used; Table S3A).

The ampliconic gene copy number-based haplotype variability observed in our data and in the data generated by Skov and colleagues (Skov et al. 2017) was mostly due to the variability of the most diverse *TSPY* and *RBMY* gene families (Fig. 8). In our data, after removing *TSPY*, the most variable gene family (Fig. 1), the total haplotype number decreased from 98 to 81. An additional removal of the *RBMY* family led to 58 haplotypes. The effect was even more dramatic for the Skov *et al.’s* data set. After removing *TSPY* from the haplotype analysis, only 19 haplotypes remained, whereas an additional removal of *RBMY* led to a significant drop to only nine haplotypes.

**Figure 8.**
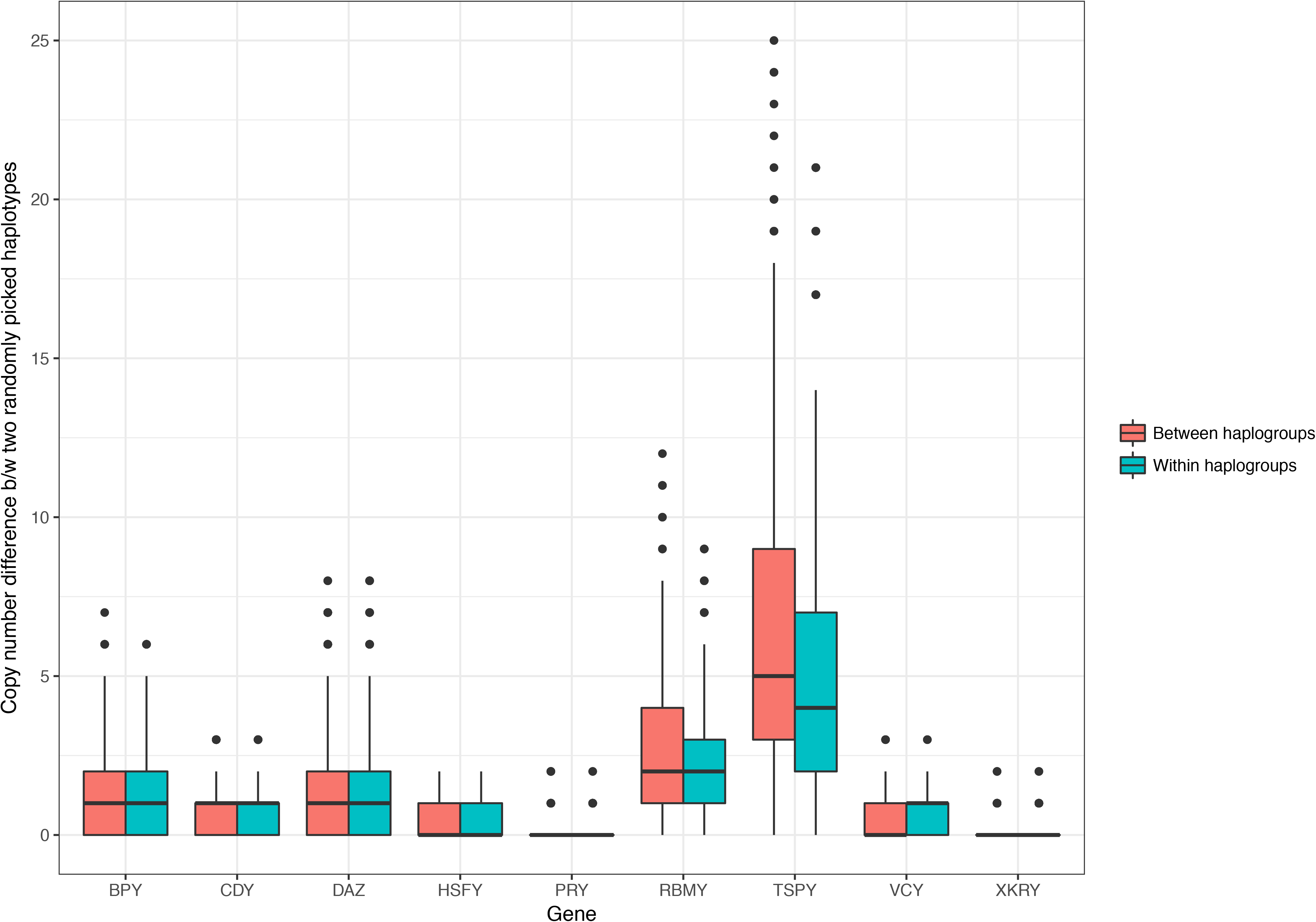
Copy number differences per ampliconic gene family between two haplotypes picked uniformly at random from within and between major Y haplogroups (1,000 samplings within and between haplogroups each; see Methods).

### Phenotypic traits

We further tested whether ampliconic gene copy number is associated with two sexually dimorphic traits, namely height and facial masculinity/femininity (FMF, see Methods). The premise here is that ampliconic genes on the Y chromosome could be involved in the development of sexually dimorphic traits. If ampliconic genes are associated with fertility, they might also have pleiotropic effects on sexually dimorphic traits. We found no statistically significant correlations between these traits and ampliconic gene copy number if we did not correct for dependence among observations due to the Y chromosomal phylogeny (Table 4). However, when we accounted for the phylogenetic relationship among Y chromosomes, height appeared to be positively correlated with copy number of *HSFY* (t-statistic = 3.272, P = 0.002), *TSPY* (t-statistic = 2.960, P = 0.004), and *XKRY* (t-statistic = 2.840, P = 0.005; Table 4). This observation suggests that people with higher copy numbers of these gene families tend to be taller. While this result is interesting, it requires further exploration. Especially important in this regard would be to study the effect of ampliconic copy number while taking into account variation in the nuclear genome.

**Table 4.**
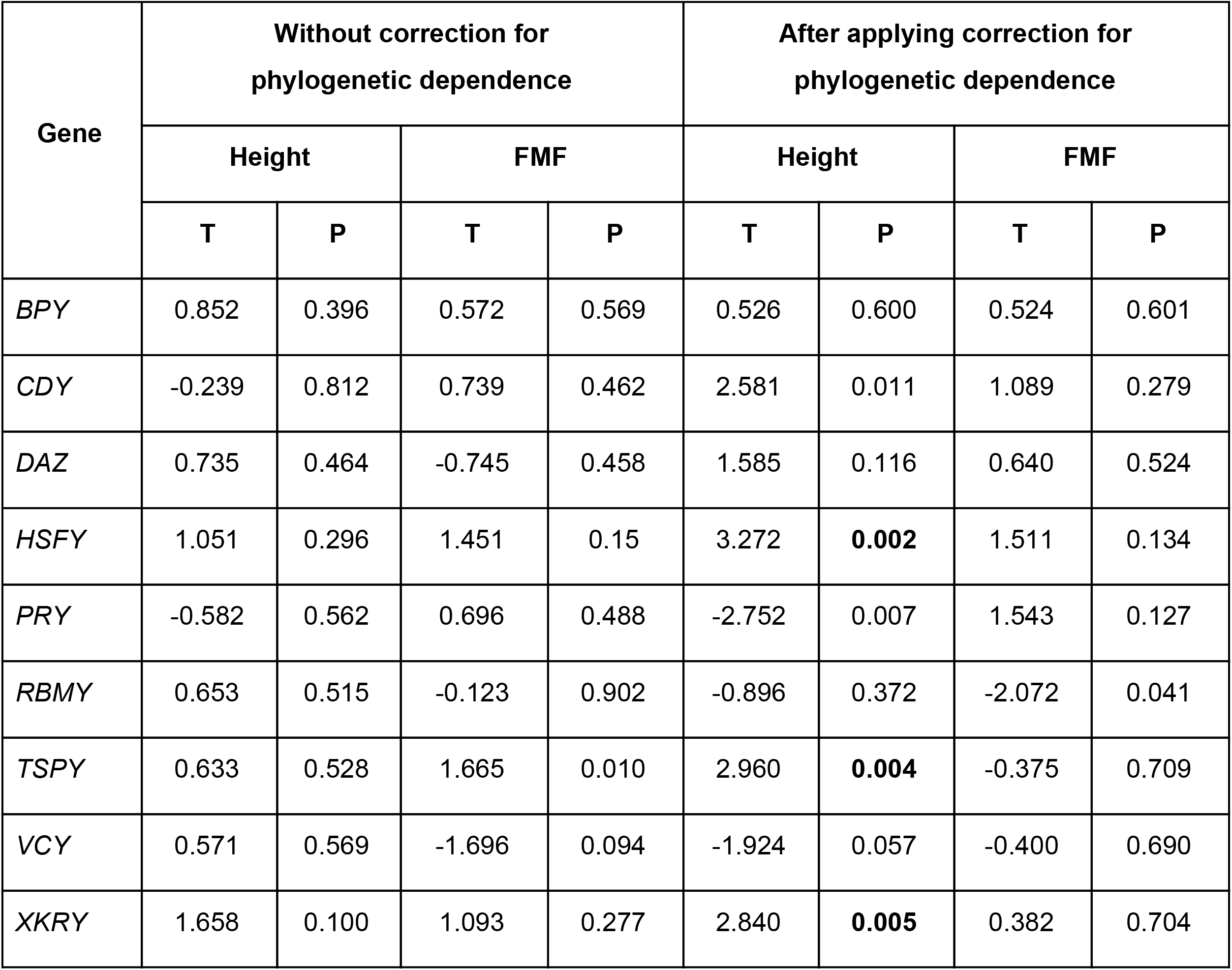
ANOVA analysis of the association between phenotypic traits (height and FMF scores) and ampliconic gene copy number without and with applying correction for phylogenetic dependence. F is the f-statistic for one-way ANOVA without correction for phylogenetic dependence. LR is the likelihood ratio between full model (predictor included) and reduced model (predictor excluded). P are the respective P values for the significance of each predictor. Significant P values after correction for multiple tests are shown in bold.

## Discussion

Very little is known about the variability in copy number of the Y chromosome ampliconic genes in humans and about how such variability impacts phenotypes. These genes, organized in nine multi-gene families, constitute 80% of only 78 protein-coding genes present on the Y chromosome (as annotated in the reference human genome) (Skaletsky et al. 2003) and are important for spermatogenesis. Here we experimentally determined the copy number of ampliconic genes in 100 individuals across the world and analyzed this variation in light of Y chromosome haplogroups based on SNPs. Additionally, we assessed whether ampliconic gene copy number is associated with two sexually dimorphic traits.

### Variability in ampliconic gene copy number

Substantial variability in ampliconic gene copy number was observed among gene families (Table 2). As a rule, gene families with high copy numbers (*RBMY* and *TSPY*) had higher variance in copy number among individuals than gene families with low copy numbers (*HSFY, PRY, VCY*, and *XKRY*). This is not surprising as the probability of gene duplication and deletion should be proportional to gene copy number, allowing for greater variation in large gene families (Ghenu et al. 2016). *TSPY* had the highest copy number and the highest level of variability from all ampliconic gene families analyzed.

In contrast to the generally low levels of nucleotide diversity on the human Y chromosome humans (e.g., (Wilson Sayres et al. 2014)), we observed high levels of variability on the Y chromosome in terms of ampliconic gene copy numbers, among individuals. A total of 98 different haplotypes were observed among 100 individuals. Thus, almost each male analyzed had his own, unique haplotype. Previously, high levels of variation in ampliconic gene copy number were reported in chimpanzee and bonobo (Oetjens et al. 2016). Thus, our results are consistent with high levels of intrachromosomal rearrangements seen on the Y chromosome (Repping et al. 2006) and with rapid evolution of Y-chromosomal multi-copy (i.e. ampliconic) genes in primates (Ghenu et al. 2016).

### Potential evolutionary mechanisms and other factors

*Mutation and drift*. Most gene families are not significantly different in their copy number among major Y chromosome haplogroups (i.e. haplogroups determined by SNPs). Only larger families – *DAZ, RBMY* and *TSPY* – showed significant differences (Table 3). In other words, most of the variation in copy number is shared among populations.

A multitude of back-and-forth duplication/deletion mutations could lead to the observed diversity of haplotypes among human world-wide populations that resulted in some homoplastic haplotypes shared by individuals belonging to different major Y haplogroups. This pattern of variation contrasts that for SNPs, which are virtually free of homoplasies and thus allow us to follow the evolution of Y chromosomes unambiguously. Interestingly, this pattern is reminiscent of that observed for microsatellite haplotype variability (Cooper 1996). Such variation patterns highlight the different nature of SNP vs. ampliconic gene copy number mutation mechanisms, but similarities between microsatellite and ampliconic gene copy number mutation mechanisms. While our purpose was not to study ampliconic gene mutational mechanisms, indirectly we can infer very rapid mutations changing ampliconic gene copy numbers that occurred among different haplotypes. More directed studies including pedigrees will have to be conducted to study the rates and relative prevalence of one-vs. multi-copy mutations in ampliconic genes from generation to generation.

*Gene conversion*. Gene conversion, prevalent at Y chromosome genes located in palindromes likely contributes to homogenization of ampliconic gene sequences, rescuing them from accumulation of deleterious mutations (Rozen et al. 2003; Betrán et al. 2012; Bellott et al. 2014). In theory, gene conversion is unlikely to influence the evolution of ampliconic gene copy number itself, because gene conversion operates at a scale smaller than individual gene copies, i. e. at the scale of a few hundreds of bases (Chen et al. 2007). Simulation studies have indicated that gene conversion acting alone does not facilitate gene duplication on the Y chromosome (Connallon & Clark 2010; Marais et al. 2010). Interestingly, it has been suggested that gene conversion can slow down the loss of redundant duplicates, nevertheless contributing to copy number evolution in this manner (Connallon & Clark 2010). Recently, gene conversion on the human Y was found to be biased towards ancestral alleles and towards GC (Skov et al. 2017). Future studies should combine sequence information of ampliconic genes together with copy number data on them to investigate Y chromosomes from humans around the globe.

*Selection*. Selection could have contributed to the observed patterns of ampliconic gene copy number variation. In particular, we observe that most of the variation in gene copy number is shared across different haplogroups. If we assume that this is not due to back mutations, uniform selection – selection that is uniform in its pressure across different human populations - could potentially explain this result (Lynch 1986; Whitlock 2008). For instance, if copy number is associated with a specific trait, and the same trait is maintained across populations by uniform selection, it might also facilitate maintenance of an optimal copy number (Hammer et al. 2008). Copy number could then be allowed to ‘drift’ around this optimum within populations by mutation.

Another potential explanation for the lack of copy number divergence across populations is balancing selection within populations via negative frequency-dependent selection (van Hooft et al. 2010). However, this contradicts the generally low nucleotide diversity on the human Y (e.g., (Dorit et al. 1995; Wilson Sayres et al. 2014) and thus is unlikely.

Our results for the comparison of between-haplogroup variation versus within-haplogroup variation based on the EVE model (Rohlfs & Nielsen 2015) suggest that the copy number of two of the nine ampliconic gene families, *TSPY* and *RBMY*, have diverged more across haplogroups than the overall level of divergence observed in all gene families together. This could be due to directional selection in one or more haplogroup lineages. However, we state this result with caution for a number of reasons. First, we only studied nine ampliconic genes and the combined pattern of divergence across these genes may not represent patterns of neutral evolution and could be skewed by one or two genes evolving non-neutrally. Second, we calculated the P values for the likelihood obtained from the EVE model assuming that the likelihood ratio follows a chi-square distribution with one degree of freedom. For the small number of genes studied here, this is a rough approximation (Rohlfs and Nielsen 2015). More sophisticated modeling is required to elucidate the role of selection on copy number in ampliconic genes.

Selection on expression levels might have also played a role in determining the observed variation in ampliconic gene copy number. Increased expression levels of some genes can lead to an increase in fitness. In this case, chromosomes carrying higher copy numbers of such genes might rise in frequency simply because a higher copy number is correlated with higher gene expression, especially for genes that are associated with fitness-related traits such as fertility (Marais et al. 2010). However, there is likely to be an upper limit for ampliconic gene copy number, as the probability of ectopic crossover events with deleterious consequences increases with the number of copies (Connallon & Clark 2010). Similarly, there might be a lower limit for each gene family, below which gene expression levels would be inadequate for spermatogenesis. These dosage-dependent factors might act as selective limits keeping copy number for ampliconic genes within a certain range (Rozen et al. 2003; Betrán et al. 2012; Bellott et al. 2014). Within this range, which might be different for each gene family, the copy number would be allowed to drift neutrally. The role of dosage-dependent selection on ampliconic gene copy number needs to be explored further by studying the relationship between ampliconic gene copy number and expression levels.

*Technical artifacts*. One potential technical factor contributing to the high haplotype variability observed for copy number variation data is amplification of pseudogenes together with functional genes. While highly accurate given the primers used, ddPCR might amplify non-functional copies if the primers anneal to them. We made a substantial effort to construct our primers in such a manner that they capture functional copies only, based on the information in the reference human chromosome Y (Tomaszkiewicz et al. 2016). However high sequence identity among gene copies might not have allowed us to completely achieve this goal. This is particularly true for the *TSPY* gene family, which is the largest tandem protein-coding array present in the human genome (Skaletsky et al. 2003). Because of its size, it is challenging to design primers that capture only functional copies of the *TSPY* family (Tomaszkiewicz et al. 2016). Other groups have reported similar difficulties with *TSPY*. For example, a recent study (Oetjens et al. 2016) used a *k*-mer based approach to detect ampliconic gene copy number variation in chimpanzees from whole-genome sequences. However, they found that the utility of their method for the repetitive *TSPY* array was limited, and their estimates of *TSPY* copy number included truncated gene copies (Oetjens et al. 2016). Ghenu and colleagues were unable to develop a robust qPCR assay to analyze *TSPY* copy number in macaques (Ghenu et al. 2016). Therefore, different methods will have to be developed to determine functional *TSPY* copy number more accurately. Nevertheless, this limitation is unlikely to be the reason behind the large number of haplotypes observed in our data. Even with the *TSPY* gene family excluded, the number of haplotypes based on ampliconic gene copy number is higher than that based on SNPs (81 vs. 39).

### Ampliconic gene copy number and male-specific sexually dimorphic traits

In this study, we tested for a potential association between ampliconic gene copy number and two sexually dimorphic traits, height and facial masculinity/femininity. We found no significant correlations between facial masculinity and copy number of any gene family. However, we detected a statistically significant positive correlation between copy number of three genes (*HSFY, TSPY*, and *XKRY*) and height. This suggests that different copy numbers of these genes might have varying downstream effects on the growth of an individual. Having said that, we state these results should be interpreted with caution for a number of reasons. Firstly, the sample size we analyzed here was relatively small (N = 100) given that the samples were taken from multiple populations worldwide. While we corrected for phylogenetic dependence among the Y chromosomes, we did not correct for variation in their nuclear genome. Sexually dimorphic traits, like many other complex traits, are likely influenced by genes located on several chromosomes. For instance, height is a polygenic trait and GWAS analyses of height have identified hundreds of common variants, each with a small effect, distributed throughout the genome (Yang et al. 2010; Wood et al. 2014). Traits specific to males and related to their reproduction are also influenced by variants located on multiple chromosomes outside of the Y. For instance, non-obstructive azoospermia, a reproductive disease characterized by the absence of sperm in semen, displays synergistic and antagonistic interactions between Y-chromosomal haplogroups and certain autosomal SNPs (Lu et al. 2016). It would be interesting to study the effect of Y ampliconic gene copy number variation on sexually dimorphic traits in light of variation in the nuclear genome.

Furthermore, future studies would benefit from focusing on males from both extremes of the trait distribution (for example, the shortest and the tallest individuals within the data set) and from the same population/haplogroup. Additionally, we only used two phenotypic traits for analysis; a more comprehensive understanding of the role of ampliconic genes and sexually dimorphic characteristics will be gained by including other traits in the analysis.

## Acknowledgements

The authors are grateful to Tomas Benjamin Gonzalez Zarzar for providing facial masculinity scores. We also thank the ADAPT study participants, without whom this research would not have been possible. Funding for the project was provided by the Penn State Center for Human Evolution and Disease (CHED) seed grant, the Huck Institutes for the Life Sciences, the Eberly College of Sciences, the Institute of Cyberscience at Penn State, and by a grant from the Pennsylvania Department of Health using Tobacco Settlement Funds. The Department specifically disclaims responsibility for any analyses, interpretations, or conclusions.

